# Cooperativity and Conformational Rearrangements in Protein-Protein and Protein-Ligand Interactions

**DOI:** 10.64898/2026.07.17.739187

**Authors:** Dinesh Thiyagaraj, Alessandro Del Re, Quan Duc Pham, Irene Gómez Garrote, Ahmad Saudi, Oleh Fedorych

## Abstract

Streptavidin-biotin, avidin-biotin interactions are classical models for protein-ligand binding, yet the energetic changes accompanying biotin binding remain poorly resolved. Using fluorescent dyes as energy sensors, we show that biotin binding produces two distinct regimes occurring in parallel as concentration of biotin increases cooperativity and conformational rearrangements, wherein cooperativity is observed via exchange broadening of fluorescence linewidth and conformational rearrangements exclusively observed in emission energy. Where the first biotin binding creates the highest contribution to the emission energy. Further analysis of tetramer-tetramer only interactions revealed extremely long ranged intermolecular interactions extending to hundreds of *nm*. The intermolecular interactions become negligible only at concentrations of approximately 10 *nM* for both streptavidin and avidin. Affinity values estimated for these diluted samples were below 1 *nM*.

## Introduction

Streptavidin and avidin are tetrameric proteins. Streptavidin-biotin and avidin-biotin interactions are among the most intensively studied interactions in molecular biology [1–5]. These interactions exhibit extraordinarily high affinity, well-defined stoichiometry and they may serve as model systems for investigations of protein-ligand interaction. After decades of research, understanding of the energetic and structural changes associated with biotin-binding events remains incomplete.

Like above, the fluorescent dye labeling has been widely used in molecular science and biology for several decades [6, 7]. Dye labels are primarily employed as optical markers that indicate the presence of molecules and their concentrations in solution. They have not been used as energy sensors capable of measuring the energy state of the labeled molecule, or more specifically, an energy value proportional to the energy of the molecule. The fluorescence linewidth and fluorescence peak position (emission energy) are two parameters that characterize the energy state of the labeled molecule. The fluorescence linewidth is related to fluorescent lifetime of the dye [8], and the fluorescence peak position is proportional to the energy of the dye. Thus, any energetic change occurring in the dye-conjugated tetramer is expected to manifest itself as a shift in the emission energy of the conjugated dye. Analysis of the dependence of the emission energy of the dye-conjugated tetramer on biotin concentration allowed us to identify two distinct contributions of biotin to the tetramer energy. It allowed us to better understand the process of occupation of tetramer sites by biotin [9–12] and revealed an unexpectedly long-range inter-tetramer interaction that cannot be explained in terms of commonly used theoretical models.

## Materials and Methods

Rhodamine 6G (Rh6G) 5-isomer was used as labelling dye and conjugated to lysine residues of avidin and streptavidin using amine-reactive crosslinking chemistry. Avidin from egg white (cat# A9275) was purchased from Sigma-Aldrich, and streptavidin (cat# 6073) from Carl Roth. Avidin and streptavidin solutions were prepared in assay buffer (100 *mM* HEPES, pH 8.0, 150 *mM* NaCl) at a final concentration of 30 µ*M* and incubated with 100 µ*M* Rh6G NHS ester, 5-isomer (cat# 24220, Lumiprobe), dissolved in 10% DMSO, in a total reaction volume of 100 µ*L*, for 15 minutes at ambient temperature, protected from light. Unreacted dye was removed using Zeba Dye and Biotin Removal Spin Columns (cat# A44296, Thermo Fisher Scientific) according to the manufacturer’s protocol. Each full spectrum spectroscopic measurement used a sample volume of 100 µ*L*. Rh6G conjugated avidin and streptavidin were assayed against biotin concentrations ranging from 10 *nM* to 1200 *nM*. Biotin (cat# B4501, Sigma-Aldrich) stock solutions, streptavidin and avidin dilutions were prepared in the assay buffer described above. To each 100 µ*L* of protein solution 0.5 µ*L* of biotin from the appropriate stock was added to achieve the desired final biotin concentration, generating a series of biotin concentrations assayed against a fixed concentration of avidin or streptavidin. In addition, the measurements were also performed in the presence of *E. coli* cells at an apparent OD600 value of 0.1, suspended in the assay buffer described above.

## Results and Discussion

As described previously, the method is based on analysis of the fluorescence linewidth and emission energy of Rh6G dye conjugated tetramers. To obtain the desired parameters, fluorescence spectra, where the fluorescence intensity is recorded as a function of wavelength (*nm*), should be recalculated on an energy scale (*eV*). All presented measurements were performed using a custom-built micro-fluorescence setup, which is described in [13, 14]. Fig. 1a) and b) show normalized fluorescence spectra measured for samples containing 200 *nM* streptavidin and 200 *nM* avidin, respectively, with increasing biotin concentration *n*_*B*_ of 0 *nM*, 10 *nM*, 50 *nM* and up to 1200 *nM*. As can be seen from the spectra, the differences between them are almost negligible and can hardly be recognized visually. Fig. 1c) and d) show the corresponding differential spectra, where the normalized spectrum measured at a given biotin concentration was subtracted from the normalized spectrum measured for the tetramer-only solution.

**FIG1.**
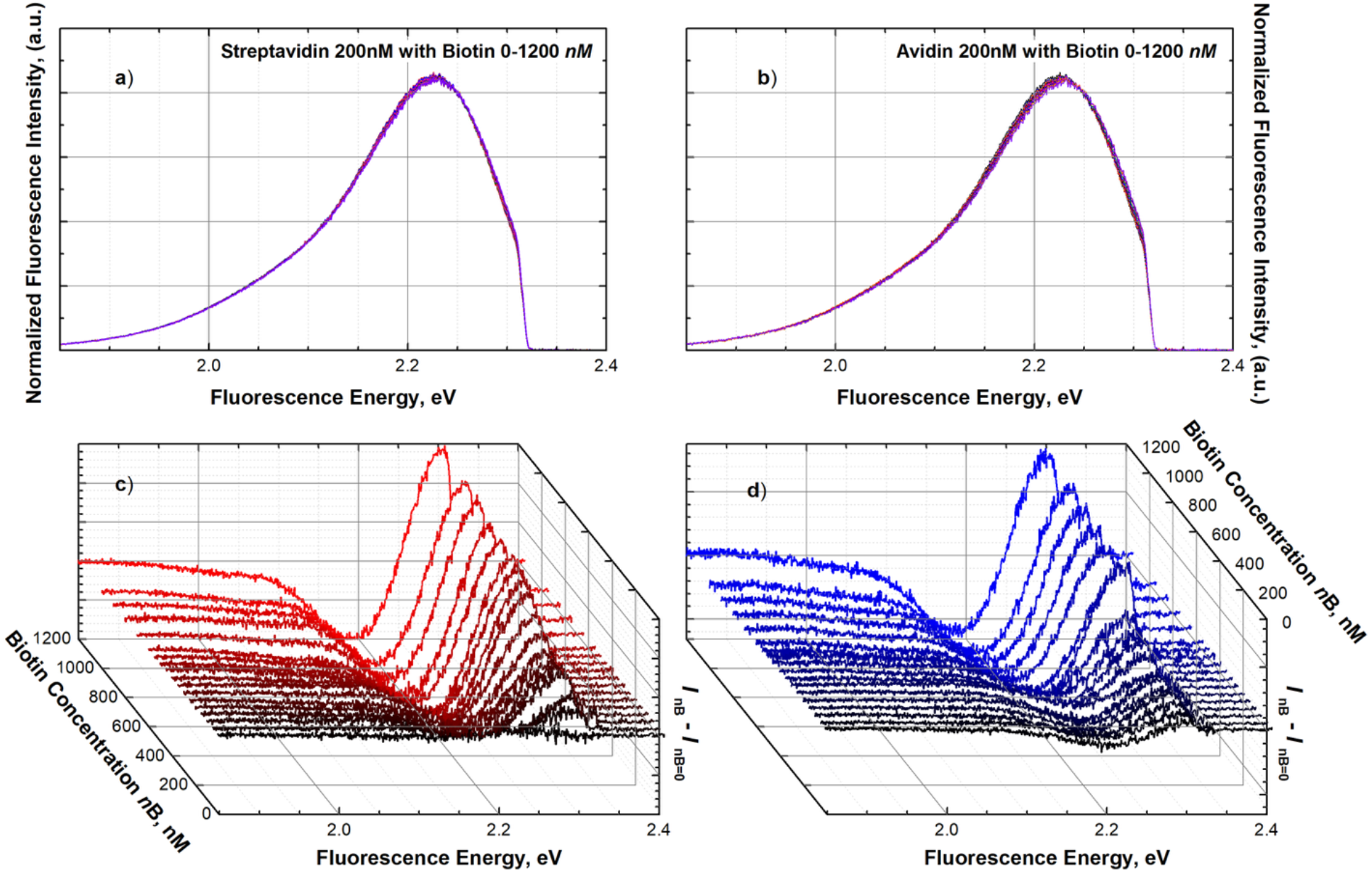
**a)** and **b)** Fluorescence spectra of 200 *nM* streptavidin and avidin respectively measured at 0 *nM*, 10 *nM*, 50 *nM* and 1200 *nM* concentration of biotin. **c)** and **d)** show the difference in normalized intensities, where the normalized spectrum without biotin was subtracted from the normalized spectrum with biotin.

The differential spectra show that increasing the biotin concentration causes a spectral shift (blue shift) of the fluorescence emission. In addition, spectra measured at higher biotin concentrations exhibit an increase in fluorescence linewidth compared with spectra obtained from tetramer-only samples. This is a very interesting result that may indicate so-called exchange broadening (due to the presence of biotin), a phenomenon well known from EPR and NMR spectroscopy [15, 16]. To investigate this effect in more detail, each fluorescence spectrum was fitted with a double-Gaussian function, and the fluorescence peak position, fluorescence linewidth, and fluorescence intensity were determined by numerical fitting. In this article, we focus on the discussion and analysis of tetramer emission energy (represented by the fluorescence peak position) and fluorescence linewidth, as well as their dependence on the concentrations of biotin and tetramers. These two parameters are less sensitive to sample purity than fluorescence intensity, where even very small changes in the refractive index of the sample buffer may lead to substantial changes in the measured fluorescence intensity. To verify the claim *E. coli* cells were added to samples containing the same tetramer concentrations. The samples containing cells were measured in the same manner as the samples without cells. The results of this investigation are presented in Fig. 2. It should be noted that, to compare samples with different tetramer concentrations, all biotin concentration dependences were normalized with respect to the tetramer concentration and plotted as dimensionless molar ratios *n*. The displaying approach simplifies the comparison of results, even when the tetramer concentration differs significantly.

**FIG2.**
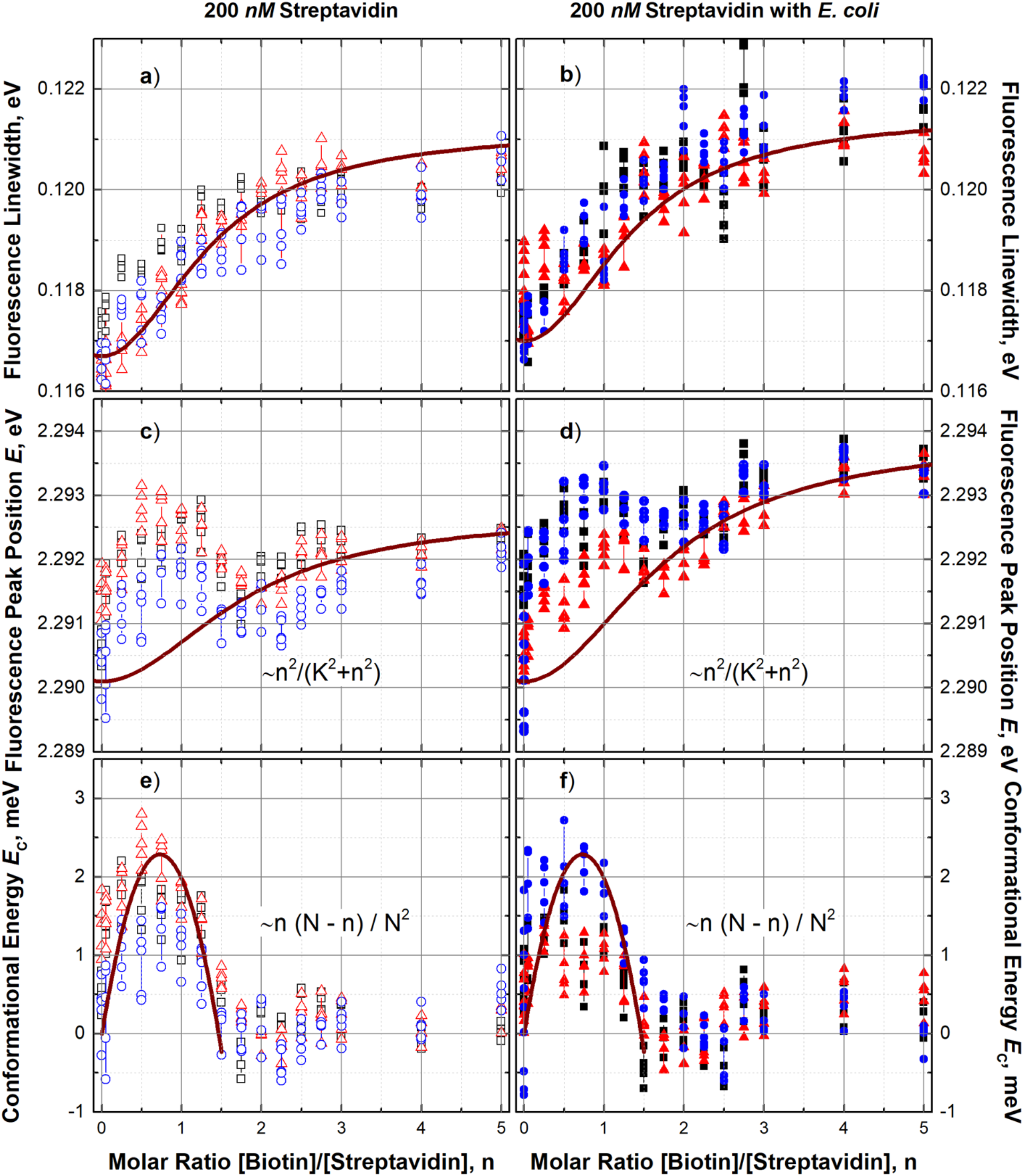
**a)** and **b)** Dependence of fluorescence linewidth on the biotin-streptavidin molar ratio for samples without (open points) and with *E. coli* cells (closed points), respectively. The brown solid line represents the theoretical fit to the experimental data using a Hill function with a Hill coefficient equal to 2. **c)** and **d)** Dependence of the fluorescence peak position on the biotin-streptavidin molar ratio (*n*) for samples without (open points) and with *E. coli* cells (closed points), respectively. The solid brown line corresponds to the theoretically predicted cooperativity between streptavidin and biotin. **e)** and **f)** Dependence of the conformational energy on the biotin-streptavidin molar ratio obtained after subtraction of the theoretically estimated cooperative contribution from the measured fluorescence peak position dependence. The extremum observed at (*n* ≈ 0.73) corresponds to conformational rearrangement of tetramer molecules induced by biotin neutralization.

An increase in biotin concentration led to an increase in the fluorescence linewidth Fig. 2a) and b). Fluorescence line broadening was observed for both types of samples, with and without *E. coli* cells. In both cases, the linewidth increased from approximately 0.117 *eV* to approximately 0.121 *eV* for biotin concentrations of 0 *nM* and 1000 *nM*, respectively, Fig. 2a) and b). The observed dependence of fluorescence linewidth on biotin concentration strongly suggests Hill-like behavior. The best agreement between the experimentally determined linewidth and the fitting model was obtained when the Hill coefficient was close to 2. This value may indicate positive cooperativity and is consistent with the behavior expected for tetramers such as streptavidin and avidin, which contain four binding sites *N* = 1,2,3,4. The dependence of fluorescence peak position on the molar ratio, as seen in Fig. 2c) and d), shows similar tendencies to those observed for the fluorescence linewidth. This behavior is particularly dominant for molar ratios greater than 2. However, at lower molar ratios, a local maximum is observed. Similar measurements were performed for other tetramer concentrations ranging from 100 *nM* to 600 *nM*. The local maximum below a molar ratio of 2 was observed for all samples, independently of their purity or tetramer type. This observation motivated us to decompose energy dependence into individual contributions, where one of the contributions should exhibit the same Hill-like dependence on biotin concentration as the previously discussed linewidth. The resulting dependence obtained after numerical subtraction of the Hill-like contribution from the energy signal is shown in Fig. 2e) and f). The energy maximum around *n* ≈ 0.73 indicates the presence of biotin associated interactions affecting streptavidin that begins immediately after the first portions of biotin were added and is completed well before reaching a molar ratio of 2.

The shape of the observed signal and its dependence on biotin concentration suggest that it could be related to the neutralization of one of the four binding sites and the associated conformational and structural modifications of the tetramer caused by occupation of a tetramer site by a single biotin molecule [10–13]. Considering four independent binding sites in each tetramer molecule, the probability of one site being occupied by biotin can be expressed as *n*/*N*. The number of possible occupation states for a tetramer molecule with *N* binding sites can be described approximately as *n*(*N* − *n*). Assuming that all sites are equivalent and that neutralization of each site produces a similar energy change, at *n* = 0 all tetramer sites are charged, while at *n* ≥ *N* all sites are neutralized. Under these assumptions, the maximum energy change should be observed at *n* = 2.

The experimental observation, however, shows that the maximum is reached at approximately *n* ≈ 0.73, showing that neutralization is not energetically equivalent for all binding sites. The best agreement between the experimental data and the model description is obtained when the energy contribution is scaled by a factor of 1/*N*^2^. This indicates that the dominant contribution to the emission energy originates from the first biotin-binding event, while the effects of subsequent biotin-binding events progressively decrease [12].

From Fig. 2, there are at least two distinct contributions to the total energy (*E*) of the tetramer molecule induced by biotin:

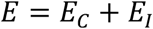

The first contribution, the biotin-induced conformational energy (*E*_*C*_), is associated with the neutralization of tetramer binding sites through their sequential occupation by biotin molecules. Its contribution to the total energy becomes negligibly small before reaching half of the molar ratio. The internal conformational rearrangement energy of the tetramer is observed only through the fluorescence peak position:

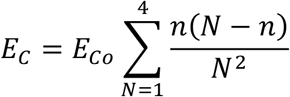

where *E*_*Co*_ ≅ 3 *meV* or *E*_*Co*_ ≅ 69 *cal*/*mole* and this value is independent of both biotin and tetramer molar ratios. This term suggests that the conformational rearrangement of a tetramer with four binding sites can be presented as a sum of four binomial contributions, with its own statistical weight. A theoretical explanation for this experimental observation is currently unknown. Nevertheless, we believe that this observation may provide useful insight into the nature of protein conformational dynamics and protein-ligand interactions.

The second contribution exhibits a Hill-like dependence on biotin concentration. The observation of this contribution in both the fluorescence peak position and, more importantly, in the increase of the fluorescence linewidth suggests that it arises from exchange broadening [15, 16] associated with cooperative interactions:

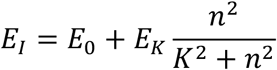

where *E*_*k*_ decreases with increasing tetramer concentration, ranging from approximately 4 − 2 *meV* or 92 − 46 *cal*/*mole*. For this type of interaction, it is important to consider that biotin acts as a coupling medium, transferring interactions and increasing spatial ordering of tetramer molecules, thereby making them more inert and slower to adapt to external perturbations. This ordering of tetramers also affects the fluorophore covalently attached to the tetramer. Stronger tetramer coupling mediated by biotin, or biotin enhanced cooperativity, results in a longer relaxation time of the tetramer, which consequently leads to a longer fluorescence lifetime of the dye. Similar behavior was observed in time-correlated measurements [17], where higher biotin concentrations resulted in longer fluorescence lifetimes of the conjugated dyes.

The observed fluorescence response cannot be described by a single binding process. Instead, the interaction of biotin with the tetramer involves an initial conformational rearrangement followed by a cooperative interaction. However, this seemingly simple and, at first sight, trivial observation has not been experimentally confirmed before. This became possible due to the ability of the analytical approach to resolve and specifically attribute the observed dependencies to their respective parameters. In particular, cooperativity appears primarily through changes in fluorescence linewidth and its broadening, while processes affecting the tetramer structure, such as conformational rearrangements, are reflected in changes of the emission energy. However, other experimental methods have addressed mainly cooperative effects.

Another conclusion, which is less scientifically significant but nevertheless has practical value, can be drawn from Fig. 2a), b), c), and d) the method is not sensitive to sample preparation and, in this specific case, to the presence of *E. coli* cells. It should be noted that if fluorescence intensity were used as the characterization parameter, it would not be possible to reliably compare or correlate even two consecutive measurements of the same sample.

While analyzing the cooperative and conformational interactions of biotin ligands with both tetramers, another observation attracted attention. The fluorescence peak position and linewidth of samples containing only tetramers increased with increasing tetramer concentration. This observation motivated a systematic study aimed at experimentally determining how fluorescence linewidth and emission energy depend on tetramer concentration. The results of this investigation are presented in Fig. 3a) and b), respectively. The tetramer concentration was converted into the average intermolecular distance, assuming initially that the molecules behave as independent entities. However, even at concentrations (distances) where tetramer molecules are expected to be isolated, measurable changes in both linewidth and fluorescence energy are observed.

**FIG3.**
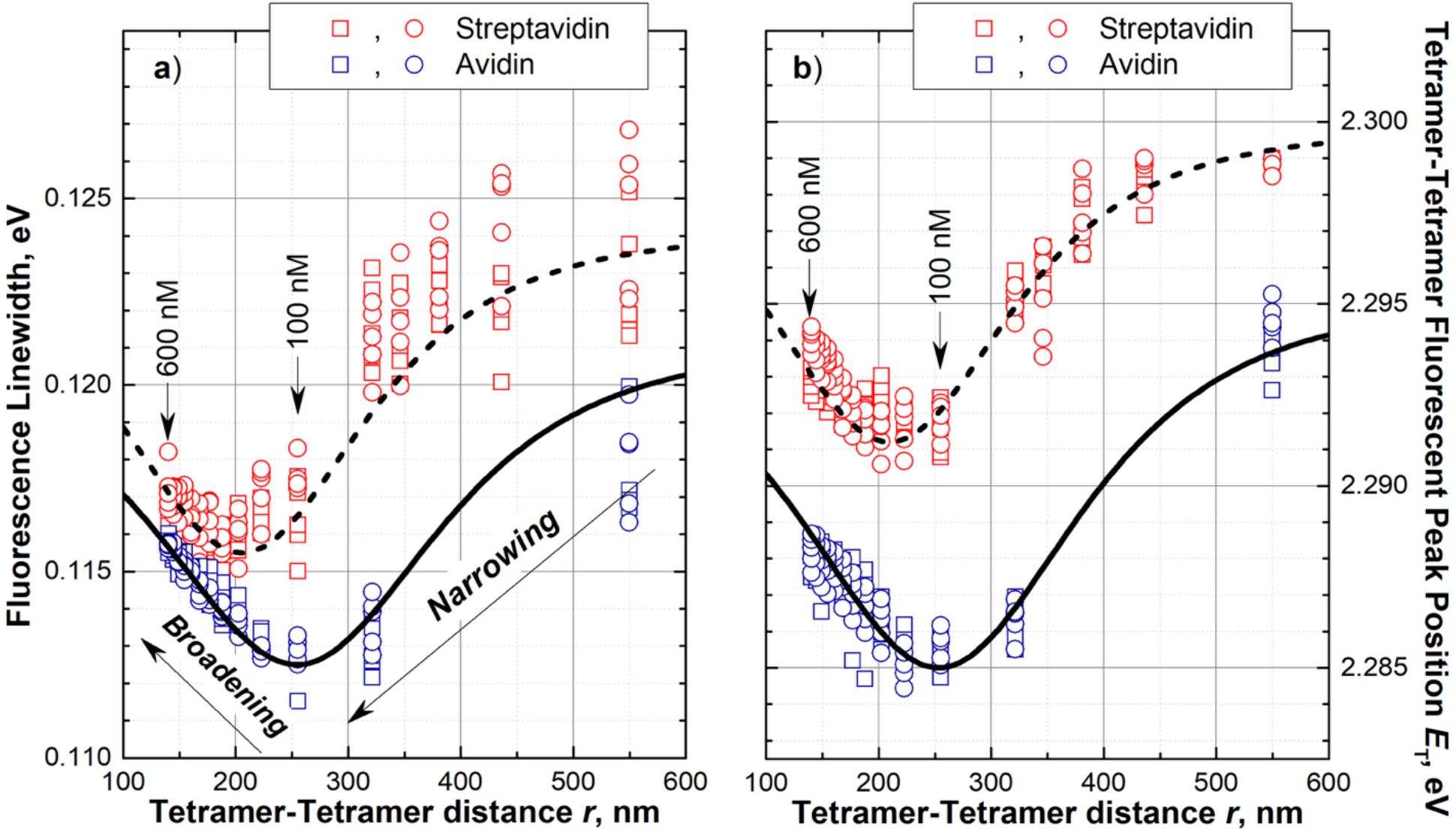
**a)** and **b)** Fluorescence linewidth and emission energy as a function of intermolecular distance, respectively. Red open points correspond to streptavidin samples, and blue open points correspond to avidin samples. The intermolecular distance was calculated from the sample concentration and used the distance dependent *E*_*T*_ equation, shown by dashed and solid black lines for streptavidin and avidin, respectively. The fluorescence linewidth **a)** initially decreases (narrowing) and subsequently increases (broadening) with decreasing intermolecular distance. The corresponding energy terms and their dependence on intermolecular distance are shown in **b)** by dashed and solid lines for streptavidin and avidin, respectively.

Initially, a decrease in linewidth and emission energy is observed up to a concentration of 200 *nM* for streptavidin (corresponding to an estimated streptavidin-streptavidin distance of approximately 200 *nm*) and approximately 100 *nM* for avidin (corresponding to an estimated avidin-avidin distance of approximately 250 *nm*). After reaching a minimum, both streptavidin and avidin exhibit an increase in fluorescence linewidth and fluorescence peak position with increasing concentration.

This is another unexpected result that may significantly affect the understanding of the nature of protein-protein interactions. At very low protein (tetramer) concentrations, approximately 10 − 20 *nM* (corresponding to intermolecular spacings of 500 − 600 *nm*), the molecules can be considered isolated and non-interacting units. Increasing the protein concentration initiates interactions between the molecules, resulting in a so-called “narrowing” effect. It should be emphasized that, in this case, no biotin or any other mediator is present that could transfer interactions between two tetramer molecules over distances of several hundred nanometers. Nevertheless, this interaction appears to reduce molecular disorder and couple the molecules, leading to a decrease in spectral width (narrowing) and, consequently, a reduction in emission energy. Further increases in tetramer concentration result in fluorescence linewidth broadening and an increase in emission energy. This broadening cannot be explained by the presence of additional components either, but rather by the decreasing intermolecular distance, which leads to stronger coupling and an increase in the effective inertia of the coupled protein molecules. There are several mechanisms that in principle could account for interactions over long ranges however, determining the specific coupling mechanism is beyond the scope of the present study. The observed effect, dependence of tetramer energy *E*_*T*_ on tetramer-tetramer distance *r*, can be described with the model [18–20]:

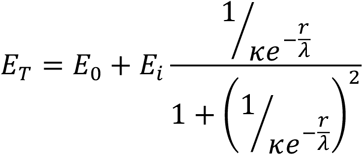

where *E*_0_ = 2.299 *eV* and 2.295 *eV, E*_*i*_ = −0.017 *eV* and −0.02 *eV, κ* = 9.1 and 10, *λ* = 95 *nm* and 110 *nm* for streptavidin and avidin, respectively. These parameters were determined by fitting the model to the experimental data.

It should be emphasized that the fluorescence peak position of Rh6G (*E*_*Rh6G*_ = 2.303 *eV*) measured in a separate experiment, is higher in energy than the *E*_0_ of streptavidin and even higher than *E*_*_ for avidin. It shows that conjugation of the tetramers shifts the Rh6G to a new emission state with different emission energy. The difference between streptavidin and avidin peak position could be related with the size of the molecules, 55 *kDa* and 66 *kDa* respectively. Larger tetramer, such as avidin, generates a stronger influence on the Rh6G, by approximately 9 *meV*, and exhibits a longer interaction range. In contrast, streptavidin changes the Rh6G energy by approximately 4 *meV* and its interaction range *λ* is shorter. This observation provides deeper insight into the method and the information it delivers. Conjugation of Rh6G to molecules, such as tetramers in this case, brings the dye into a new energetic state determined by the influence of the tetramer on the dye. All subsequent structural and energetic changes that the conjugated tetramer undergoes will be reflected by the dye. However, these changes should not be interpreted as absolute energies of the tetramer itself, but rather as relative changes with respect to the initial state of the conjugated dye-tetramer system, where *E*_*T*_ = *E*_0_.

As shown in Fig. 3, at tetramer concentrations above approximately 20 *nM*, the tetramer molecules can no longer be considered isolated, and tetramer-tetramer interactions become measurable. Tetramer-tetramer interactions will also affect the tetramer-biotin binding affinity. On the other hand, samples with low tetramer concentrations, e.g. 10 *nM*, emit significantly lower fluorescence intensities, resulting in a low signal-to-noise ratio and increased variance of the determined parameters (linewidth and emission energy). Nevertheless, experiments involving increasing biotin concentration were performed using samples containing 10 *nM* tetramer, both streptavidin and avidin. The biotin concentration was increased from 0 to 10 *nM* and subsequently in 50 *nM* increments. As in the previous experiments, the recorded spectra were analyzed, and the fluorescence linewidth and fluorescence peak position were determined for each spectrum.

The fluorescence linewidth and fluorescence peak position as functions of the biotin-tetramer molar ratio *n* are shown in Fig. 4a) and b). As can be seen from the figure, even the initial addition of 10 *nM* biotin triggers a measurable tetramer-biotin interaction, and by 50 *nM* the effect has already reached saturation. As observed, the interaction leads to a reduction in energy, in contrast to the behavior measured for tetramer concentrations above 50 *nM*, see Fig. 2a), b), c) and d), where the energy increased with increasing concentrations of biotin. A decrease in fluorescence peak position is clearly observed for streptavidin and avidin, see Fig. 4b). Modeling of these observations suggests that the affinity of both tetramers is below 10 *nM* and is likely below 1 *nM*.

**FIG4.**
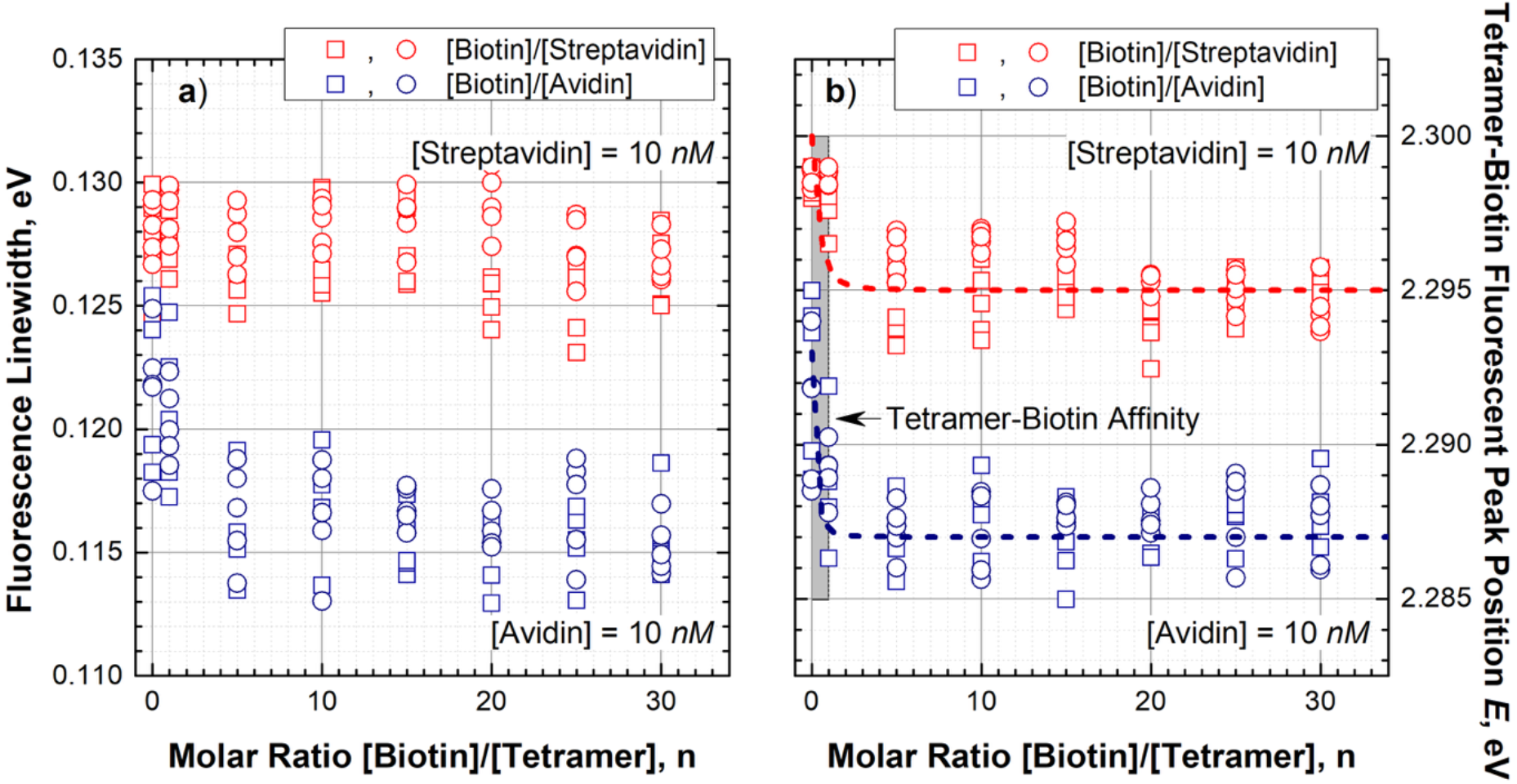
**a)** and **b)** Fluorescence linewidth and tetramer-biotin fluorescent peak position as a function of the tetramer-biotin molar ratio, respectively. The concentrations of streptavidin (open red points) and avidin (open blue points) were equal to 10 *nM*. Dashed red and blue lines show the theoretical dependence of the interaction energy described by a Hill function with a Hill coefficient equal to 2 and an affinity of less than 1 *nM* is shown by the gray region.

## Conclusions

It was demonstrated that fluorescent dye labeling of protein molecules allows the use of the fluorescence peak position as the value revealing the energy transformations of the molecule. This parameter makes the measurements less sensitive to the sample purity, including, in our case, the intentional addition of *E. coli* cells. Analysis of the fluorescence peak position in combination with fluorescence linewidth made possible the observation of conformational effects associated with the neutralization of tetramer binding sites by biotin. It was demonstrated that the strongest conformational effect is associated with the binding of the first biotin molecule. Furthermore, we were able to determine that tetramers exhibit interaction ranges extending over hundreds of nanometers. In the concentration range where tetramer-tetramer interactions are negligible, we showed that biotin binding to an isolated tetramer leads to a reduction in the emission energy, with an estimated binding affinity stronger than 1 *nM*.

## Acknowledgments

We thank to the Investitionsbank Berlin (IBB) for the financial support.

## Author Contributions

Conceptualization and Investigation: Dinesh Thiyagaraj, Oleh Fedorych.

Methodology, Formal Analysis, Visualization: Oleh Fedorych.

Writing – original draft: Oleh Fedorych.

Writing – review & editing: Dinesh Thiyagaraj, Oleh Fedorych, Alessandro Del Re. Resources – Duc Quan Pham, Irene Gómez Garrote, Ahmad Saudi

