## Supplementary File1 for "Cooperativity and Conformational Rearrangements in Protein-Protein and Protein-Ligand Interactions"

### Supporting information

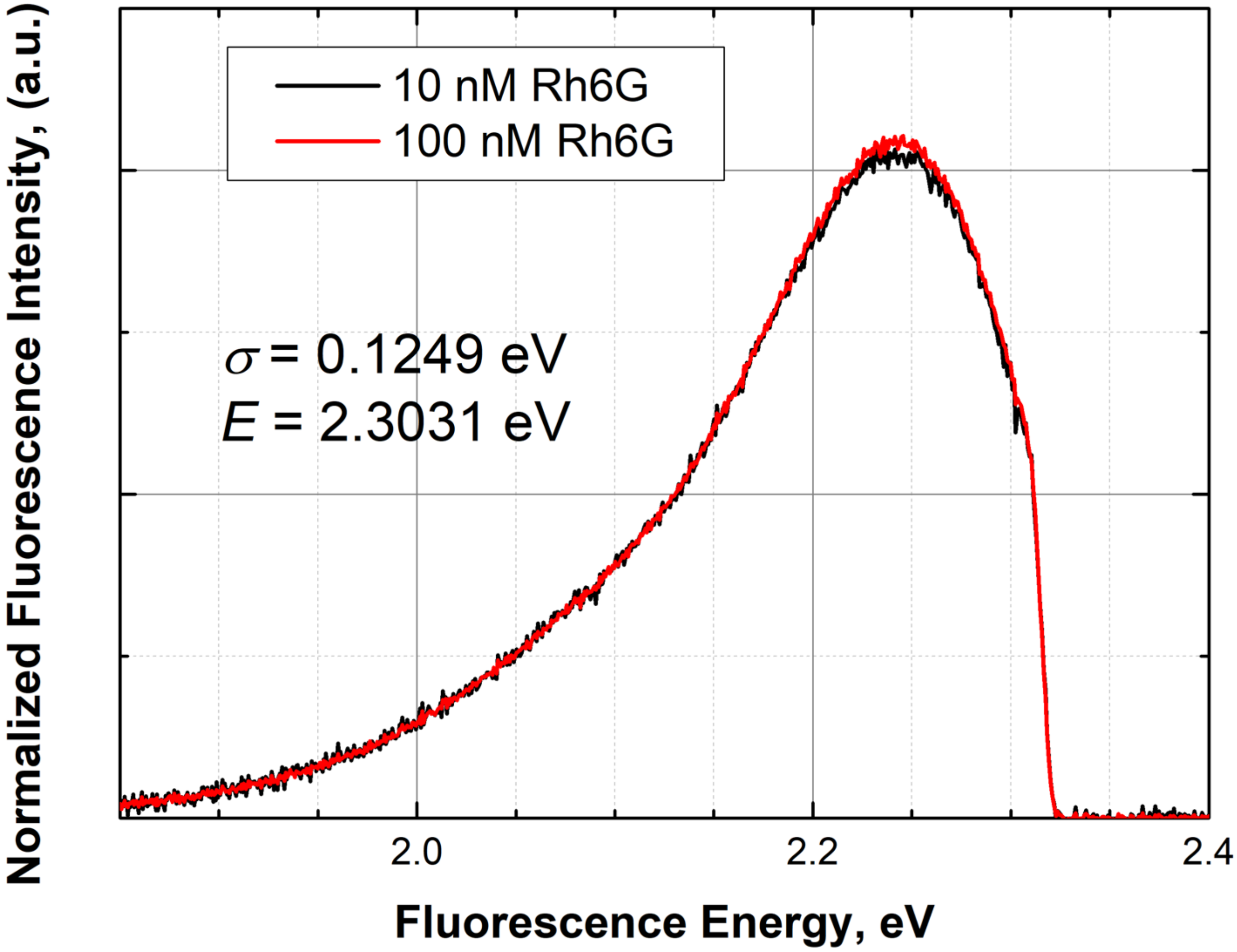


**FIG.S1**. Normalized fluorescence spectra of $10\,nM$ (black line) and $100\,nM$ (red line) Rhodamine 6G in assay buffer. Both spectra exhibit the same fluorescence linewidth $\sigma = 0.1249\ eV$ and fluorescence peak position $E = 2.3031\ eV$.
